# Identification of a functional non-coding variant in the GABA_A_ receptor α2 subunit of the C57BL/6J mouse reference genome: Major implications for neuroscience research

**DOI:** 10.1101/540211

**Authors:** Megan K. Mulligan, Timothy Abreo, Sarah M. Neuner, Cory Parks, Christine E. Watkins, M. Trevor Houseal, Thomas M. Shapaker, Michael Hook, Haiyan Tan, Xusheng Wang, Jesse Ingels, Junmin Peng, Lu Lu, Catherine C. Kaczorowski, Camron D. Bryant, Gregg E. Homanics, Robert W. Williams

## Abstract

GABA type-A (GABA-A) receptors containing the α2 subunit (*Gabra2*) are expressed in most brain regions and are critical in modulating inhibitory synaptic function. Genetic variation at the GABRA2 locus has been implicated in epilepsy, affective and psychiatric disorders, alcoholism and drug abuse. *Gabra2* expression varies as a function of genotype and is modulated by sequence variants in several brain structures and populations, including F2 crosses originating from C57BL/6J (B6J) and the BXD recombinant inbred family derived from B6J and DBA/2J. Here we demonstrate a global reduction of *Gabra2* brain mRNA and protein in the B6J strain relative to other inbred strains, and identify and validate the causal mutation in B6J. The mutation is a single base pair intronic deletion located adjacent to a splice acceptor site that only occurs in the B6J reference genome. The deletion became fixed in B6J between 1976 and 1991 and is now pervasive in many engineered lines, BXD strains generated after 1991, the Collaborative Cross, and the majority of consomic lines. Repair of the deletion using CRISPR-*Cas9* mediated gene editing on a B6J genetic background completely restored brain levels of *Gabra2* mRNA and protein. Comparison of transcript expression in hippocampus, cortex, and striatum between B6J and repaired genotypes revealed alterations in GABA-A receptor subunit expression, especially in striatum. These results suggest that naturally occurring variation in GABRA2 levels between B6J and other substrains or inbred strains may also explain strain differences in anxiety-like or alcohol and drug response traits related to striatal function. Characterization of the B6J private mutation in the *Gabra2* gene is of critical importance to molecular genetic studies in neurobiological research as this strain is widely used to generate genetically engineered mice and murine genetic populations, and is the most widely utilized strain for evaluation of anxiety-like, depression-like, pain, epilepsy, and drug response traits that may be partly modulated by *Gabra2* function.

## Introduction

GABA-A receptors are pentameric ligand gated chloride channels responsible for fast inhibitory neurotransmission. The sequence, structure, and chromosomal location of receptors and their cognate subunits are highly conserved among mammals (Olsen and Sieghart, 2009). Receptors are typically composed of two α, two β, and a γ or δ subunit with the most common receptor subtypes containing α1 or α2 subunits. Subunit composition can alter receptor pharmacology and localization (Sieghart, 1995). Receptors containing the α2 subunit (*Gabra2*) are abundantly expressed in mouse hippocampus, frontal cortex, amygdala, dorsal and ventral striatum, and hypothalamus—brain regions important for motivation, reward, anxiety, depression, and fear. Over 500 common human variants have been identified at the *GABRA2* locus, including numerous intronic variants, a handful localized to the UTRs, and one synonymous variant in exon 4. Genetic variation at this locus has been linked to alcohol dependence (Covault et al., 2004; Edenberg et al., 2004; Lappalainen et al., 2005; Agrawal et al., 2006; Enoch et al., 2006; Fehr et al., 2006; Lind et al., 2008a; Lind et al., 2008b; Soyka et al., 2008; Villafuerte et al., 2013; Li et al., 2014), the subjective effect of alcohol (Pierucci-Lagha et al., 2005; Roh et al., 2011; Uhart et al., 2013; Arias et al., 2014), excess EEG fast activity (Lydall et al., 2011), cocaine reward (Dixon et al., 2010a), substance dependence (Enoch et al., 2010), epilepsy (International League Against Epilepsy Consortium on Complex Epilepsies. Electronic address, 2014), and impulsivity and insula cortex activity during reward or loss anticipation (Villafuerte et al., 2013). Furthermore, interactions between early life stress and *GABRA2* polymorphisms may influence vulnerability to addiction (Enoch, 2008; Dick et al., 2009; Enoch et al., 2010). However, the impact of these variants on expression, isoform usage, or function of GABRA2 mRNA or protein is unclear (Myers et al., 2007; Heinzen et al., 2008; Gibbs et al., 2010; Liu et al., 2010; Kim et al., 2014; Lieberman et al., 2015).

Preclinical models can be useful in dissecting enigmatic relationships between human behavior and genetic variation. Genetically engineered mouse models in which *Gabra2* expression or sensitivity to modulation has been altered (knock-out and knock-in) suggest a role for GABA-A receptors containing the α2 subunit in depression (Vollenweider et al., 2011), alcohol intake and behavioral response to alcohol (Boehm et al., 2004; Blednov et al., 2011; Liu et al., 2011; Blednov et al., 2013), anxiety (Dixon et al., 2008), and cocaine-conditioned reinforcement and behavioral sensitization (Morris et al., 2008; Dixon et al., 2010b). Genetic deletion of *Gabra2* is also associated with a decrease in miniature inhibitory postsynaptic current in the nucleus accumbens core (Dixon et al., 2010b), suggesting an important role of *Gabra2* in modulating inhibitory signaling in this region. However, no studies to date have characterized naturally occurring variants segregating among murine populations that could alter *Gabra2* levels or function, and provide insight into the role of human variants.

In this study, we leverage naturally occurring variation in murine genetic populations of varying complexity in order to better understand the role of *Gabra2* variants on gene function. We previously profiled gene expression data from multiple mouse strains and crosses, including nearly isogenic C57BL/6 substrains and the well-characterized BXD family of strains derived from a cross between B6J and DBA/2J (D2) inbred mice, to characterize genetic variation in *Gabra2* expression (Mulligan et al., 2012). Here we identify the source of this variation as a spontaneous intronic deletion in B6J that underlies variation in *Gabra2* mRNA and protein expression across diverse murine populations that incorporate B6J as a progenitor strain. We have quantified molecular and behavioral consequences of the private mutation in B6J by CRISPR-*Cas9* genome editing. This work demonstrates the functionality of often ignored non-coding variants in the mouse genome and characterizes the downstream consequences of *Gabra2* variation on molecular traits.

## Methods

### Mice

C57BL/6J-*Tyr*<*c-2J*> mice were purchased from Jackson Laboratories and C57BL/6NCrl mice were purchased from Charles River Laboratories at 8 weeks of age and allowed to acclimate for 9 days at UTHSC before tissue collection. C57BL/6J, DBA/2J, C57BL/6NJ, C57BL/6JEiJ, and C57BL/6ByJ are maintained in-house up to generation 6 at UTHSC from Jackson Laboratory stock. BXD mice are maintained at UTHSC. The B6-*Gabra2*^em1Geh/^J CRISPR repair line was generated at the University of Pittsburgh (as described below) and is maintained as a heterozygous breeding colony by the Mulligan laboratory at UTHSC. All mice were housed in same-sex cages, allowed a*d-lib* access to food and water and maintained on a 12h/12h light/dark cycle in a climate-controlled facility. Only adult mice (between 70 and 200 days of age) were included in the study. All animal care and handling procedures were approved by the UTHSC and the University of Pittsburgh Institutional Animal Care and Use Committees.

### Data sets used

The following data sets available at GeneNetwork.org were used in the analysis: GN76 OHSU/VA B6D2F2 Brain mRNA M430 (Aug05) RMA (Hitzemann et al., 2004), GN84 OHSU/VA B6D2F2 Striatum M430v2 (Sep05) RMA (Hitzemann et al., 2004), GN281 INIA Hypothalamus Affy MoGene 1.0 ST (Nov10) (Mozhui et al., 2012), GN323 INIA Amygdala Cohort Affy MoGene 1.0 ST (Mar11) RMA, GN110 Hippocampus Consortium M430v2 (Jun06) RMA (Overall et al., 2009), GN175 UCLA BHHBF2 Brain (2005) mlratio (Orozco et al., 2009; van Nas et al., 2010), GN171 UCLA CTB6/B6CTF2 Brain (2005), mlratio, INIA LCM (11 Regions) BASELINE RNA-seq Transcript Level, GN206 UMUTAffy Hippocampus Exon (Feb09) RMA, and GN732 GTEXv5 Human Brain Hippocampus RefSeq (Sep15) RPKM log2 (Consortium, 2013). In addition to references provided, additional information on these data sets can also be found at GeneNetwork.org.

### Tissue collection

The cortex, hippocampus and striatum (dorsal and ventral) were dissected from up to three males and three females for each strain (effective strain *n* = 5 or 6). At time of tissue harvest, purchased mice were ~9 weeks and in-house strains were ~8 weeks of age. Tissue was immediately frozen in liquid nitrogen and stored at –80 °C until further use. Additional peripheral tissue, including the liver and spleen, was also dissected from each strain and stored at –80 °C until further use. Whole brains from wild-type *Gabra*2^B6J/B6J^, homozygous *Gabra2*^KI/KI^ and heterozygous *Gabra2*^KI/-^ mice were dissected and placed in glass scintillation vials, which were subsequently kept at –80°C. To prepare for sub dissection of whole frozen brains, the brains were kept at –20 °C at least 30 min prior to dissection. Whole brains were placed dorsal side down into a brain matrix, with two coronal cuts being made by single edge razor blades to obtain striatal tissue. The first cut was made rostral and proximal to the optic chiasm (bregma –3.0), and the second cut was made two millimeters forward from the first cut (bregma +2.0). Hippocampus and cortex were then sub-dissected from the brain tissue remaining caudal to the first razor blade cut. *Gabra2* homozygous knockout brains were kindly provided by Y. Blednov at the University of Texas at Austin. These mice were maintained on a mixed C57BL6Jx129/SvEv background at the University of Sussex and were backcrossed twice to B6J at the University of Texas. Tissue was flash frozen in liquid nitrogen, and immediately (within 1–2 days) shipped on dry ice to UTHSC where cortex, hippocampus and striatum were dissected and stored at –80 °C until further use.

### qPCR to validate mRNA levels

Five biological replicates were used for each strain and consisted of both males and females (two to three of each sex). RNA was isolated from brain tissue using the RNeasy Lipid Tissue kit (Qiagen), with cDNA then being generated from 1 μg of RNA utilizing the First Strand Reverse Transcriptase kit (Roche). Quantitative PCR was utilized to assess target gene expression using the KAPA SYBR Fast QPCR Master Mix, probes from Roche’s Universal ProbeLibrary, custom primers from Integrated DNA Technologies (IDT), and a Roche Light Cycler 480 II. Universal probes and custom primers were selected and designed, respectively, using the Universal Probe Library (www.universalprobelibrary.com; Roche Diagnostics). For *Gabra2* (NM_008066.3) we used Universal Probe #103 and the following primers: Left–ACAAAAAGAGGATGGGCTTG and Right–TCATGACGGAGCCTTTCTCT. Cyclophilin D (NM_026352.3) was selected as the endogenous control gene for all tissues (Schmittgen and Livak, 2008). Data was analyzed using the ΔΔCt method as described in (Livak and Schmittgen, 2001), with C57BL/6J selected as the reference or calibrator strain to calculate fold-change expression. For gene expression in the CRISPR mice, the coefficient of variation was quantified using the ΔCt method. All statistical tests were conducted in R using R Studio (RStudio: Integrated Development for R. RStudio, Inc., Boston, MA). One-way ANOVA was performed for comparing more than two independent samples, with Tukey’s HSD post-hoc test subsequently applied to assess pairwise significance between genotypes.

### Validation of genomic variants

Variants that were polymorphic between the B6J reference genome and B6NJ and D2 were identified using 100X coverage of the D2 genome available at http://ucscbrowser.genenetwork.org/ (Wang et al., 2016) and the Welcome Trust Sanger Institute (sequenced genomes available for 36 inbred mouse strains, including the B6NJ substrain, at http://www.sanger.ac.uk/cgi-bin/modelorgs/mousegenomes/snps.pl). Initially we identified two high quality indels located on Chr 5 at 71,014,638 bp (deletion) and 71,031,384 bp (rs225241970, deletion), and a highly conserved intergenic SNP located at 70,931,531 bp (all coordinates given based on the mm10 assembly). Genomic DNA was extracted from B6J, D2, B6NJ, B6EiJ, B6ByJ, BXD29, and BXD40 using the DNeasy Blood & Tissue Kit (Qiagen) from frozen liver or spleen tissue. Variants were genotyped using primers designed against the reference genome using the Primer3Plus web tool and then purchased from IDT. Primers targeting each variant include: (1) SNP rs29547790@70,931,531 Mb Left–AAAAGTCAGGGTGTGGTTGG and Right–GGAGTGCAGCTCTCTCTTTTGG, (2) indel@71,014,638 Mb Left–TCAGGAGTCCAGATTTTGCTG and Right–TCTCTCAGTTCCGTTTTCTGTAA, and (3) indel rs225241970@71,031,384 Mb Left–AGCACCCTTGGGAAGAAAGG and Right–GGTCTCATCAGGAAATAGAACCGA. All coordinates correspond to the mouse GRCm38/mm10 assembly. Genomic intervals (~100–300 bp) containing the variants were amplified by PCR and unincorporated nucleotides and primers removed using ExoSAP-IT (Thermo Fisher Scientific). Traditional Sanger capillary sequencing was performed by the UTHSC Molecular Resource Center Institutional Core using the ABI Prism 3130 Genetic Analyzer system. The only variant that could not be confirmed was the indel rs225241970 located on Chr5 at 71,031,384 bp as none of the seven strains tested, including B6J, were polymorphic at this position. This genomic region includes a microsatellite repeat (CA)n that may have led to sequencing errors in the reference genome.

### Traditional western analysis

Lysates were prepared from frozen tissue and protein concentration was determined using a Nanodrop2000 Spectrophotometer (ThermoScientific). 50 μg of total protein was loaded and separated on a 7.5% SDS-PAGE gel. Proteins were transferred using the Bio-Rad TurboTransfer system and blocked for 30 minutes at room temperature. For chemiluminesence western blots, blots were incubated in primary antibody for GABRA2 (1:500 rabbit polyclonal, PhosphoSolutions, Cat #822-GA2CL) overnight at 4**°**C. Blots were washed 3×5mins with blocking buffer and incubated with horseradish-peroxidase conjugated antibody for 1h at room temperature. Blots were developed using the Super Signal West Pico Chemiluminescent Substrate kit (ThermoScientific #34080) on a BioRad ChemiDoc imaging system. Blots were then incubated in a stripping solution (Restore Western Blot Stripping Buffer, ThermoScientific #21059) for 8 minutes, re-probed overnight at 4C with a GAPDH antibody (1:5000, mouse monoclonal, Fitzgerald, Cat #10R-G109A), and developed again the following day. For fluorescent western blots, blots were incubated with both primary antibodies together overnight at 4C, followed by anti-rabbit and anti-mouse fluorescent-conjugated antibodies. Visualization was performed using an Odyssey image scanner. Blot intensities across all blots were quantified using ImageJ. Original blots can be found in SFigs 1 and 2.

**Fig. 1.**
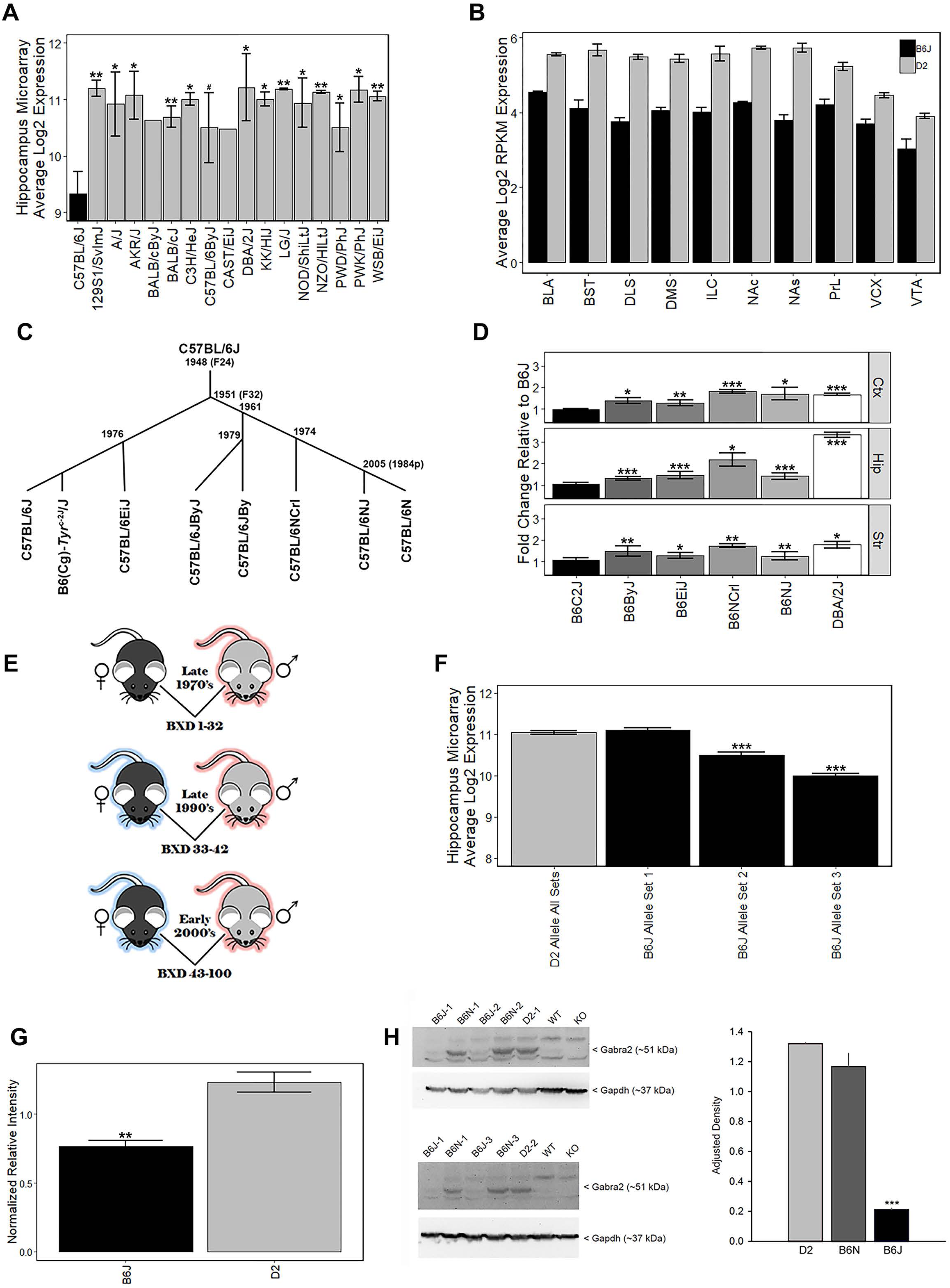
Strain variation in *Gabra2* expression. (A) Low expression of *Gabra2* in B6J brain relative to other inbred strains and substrains. B6J (blue) showed the lowest expression of *Gabra2* in hippocampus (data set GN110, probe set 1421738_at) compared to 15 inbred strains, including the D2 strain (red), and a closely related B6 substrain (grey). (B) *Gabra2* levels measured by RNAseq are reduced in 10 mesocorticolimbic regions relative to D2, indicating a global reduction in B6J. BLA = basolateral amygdala, BST = bed nucleus stria terminalis, DLS = dorsolateral striatum, DMS = dorsomedial striatum, ILC = infralimbic cortex, NAc = nucleus accumbens core, NAs = nucleus accumbens shell, PrL = prelimbic cortex, VCX = visual cortex, VTA = ventral tegmented area. All *P*s < 0.001 for within region contrasts between B6J and D2. (C) Approximate time line seperating C57BL/6 substrains. (D) Average *Gabra2* expression level (measurd by qPCR) in the cortex, hippocampus, and striatum of D2, C57BL/6 substrains, and congenic strains relative to B6J shown at left. The B6N lineage (seperated in 1951 at generation F32 from B6J) includes C57BL/6JByJ (B6ByJ), C57BL/6NCrl (B6NCrl), and C57BL/6NJ (B6NJ) and all showed higher expression of *Gabra2* relative to B6J. Even the C57BL/6EiJ (B6EiJ) substrain that diverged most recently from C57BL/6J in 1976 showed higher brain expression of *Gabra2*. The only substrain with low *Gabra2* expression similar to C57BL/6J is the B6(Cg)-*Tyr*^c-2J^/J (B6C2J) congenic strain. This albino mutation was detected in 1970 but the original strain harboring the spontaneous mutation was backcrossed to B6J creating a congenic line. All statistical contrasts performed relative to B6J. (E) Derivation of BXD strain cohorts. Each cohort was derived by separate crossing and inbreeding of female B6J and male D2 parental inbred strains. The first cohort of BXD strains (1 through 32) was derived in the late 1970s. Another set of strains (33 through 42) was produced in the early 1990s. The last cohort of BXD strains (43 to 100) was created in the early 2000s. (F) Hippocampal differences in *Gabra2* expression between early and later derived BXD strains. There was little difference in the expression of *Gabra2* between BXD strains with the B6 (black) or D2 allele (red) in the first cohort. Hippocampal differences in *Gabra2* expression between early and later derived BXD strains. There was a wide range of variation in *Gabra2* expression between inheritance of B6 and D2 alleles in the BXD population. While there was little variation in *Gabra2* expression associated with inheritance of the D2 allele, there was a large amount of within genotype variation for the B6 allele. However, inheritance of the B6 allele in the later two cohorts was associated with a dramatic reduction in *Gabra2* expression. This suggests the appearance of a mutant allele in the B6J line that occurred after the late 1970’s and prior to 1990. Data from GeneNetwork data set GN110, probe set 1421738_at. (G) Protein levels of *Gabra2* were reduced in B6J relative to D2 using an unbiased shotgun proteomics approach. Relative expression of *Gabra2* protein shown as normalized relative intensity for *Gabra2* in B6J and D2 hipppocampus. Eight different peptide sequences that matched *Gabra2* were used to generate normalized relative intensity counts for each sample. Each sample is 3 pooled animals (males only). (H) Hippocampal protein levels of *Gabra2* are reduced in B6J (n = 3) compared to the closely related B6N (n = 3) substrain and D2 (n = 2). Western blot is shown to the left and relative density shown to the right. The experiment was repeated twice and both B6J sample 1 and B6N sample 1 were run each time. Tissue from *Gabra2* KO mice and their wild type control strain (WT) were kindly provided by Y. Blednov at The University of Texas. These mice were originally maintained on a mixed B6J and 129/SvEv background but were backcrossed twice to B6J upon arrival at The University of Texas, thus the WT control strain has low *Gabra2* protein expression similar to B6J. ANOVA for effect of strain is significant F(2,5) = 103.9, p < 0.001. Post-hoc tests show B6J is different from both B6N and D2 (p < 0.001) but B6N and D2 are not significantly different from each other (p = 0.15). Significance defined as: * = *p* < 0.05, ** = *p* < 0.01, *** = *p* < 0.001.Significance defined as: # = *p* < 0.1, * = *p* < 0.05, ** = *p* < 0.01, *** = *p* < 0.001.

**Fig. 2.**
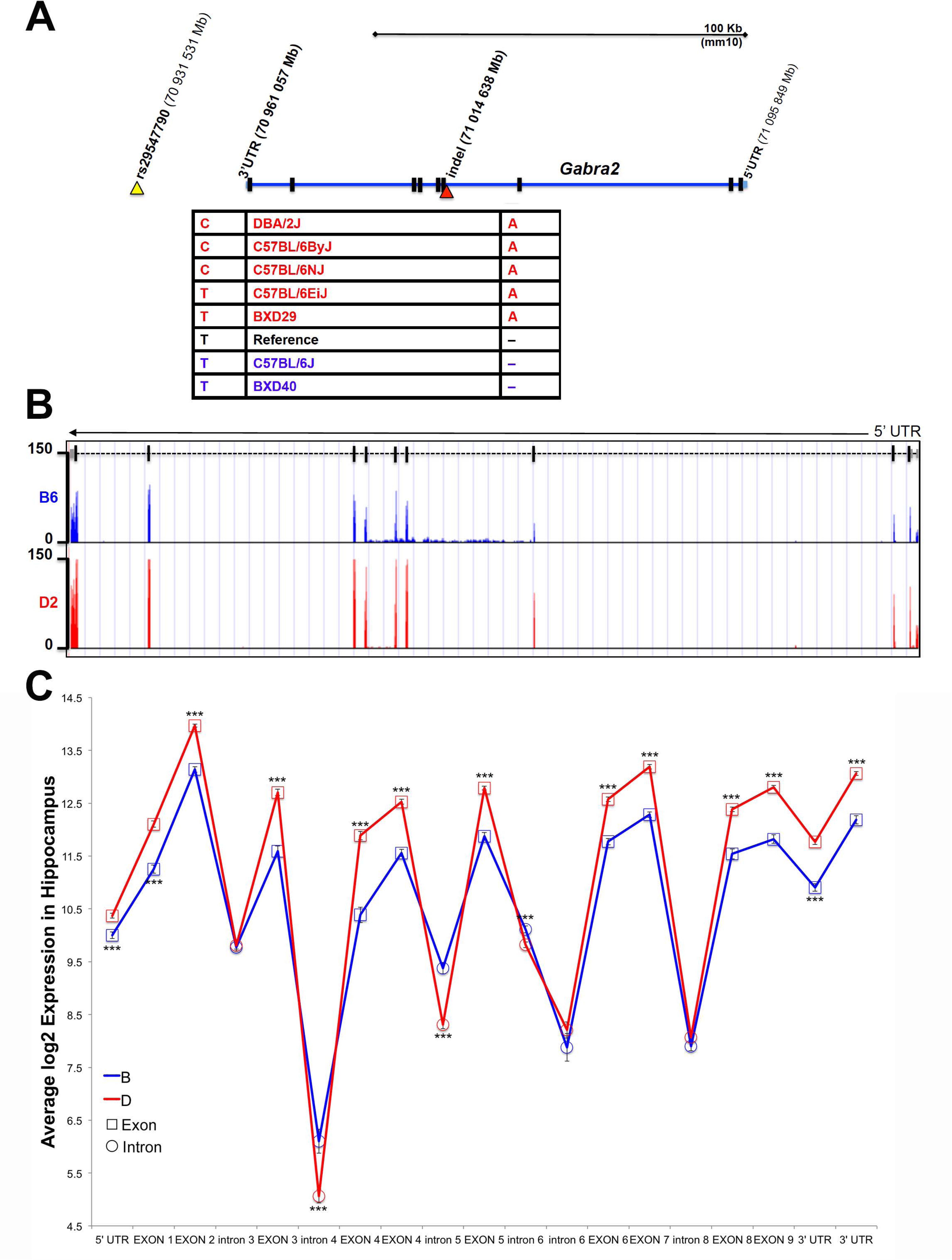
Identifcation of private variant in *Gabra2*. (A) Position of variants near the *Gabra2* gene locus. An intergenic SNP (yellow arrohead) and intronic indel (red arrowhead) were the only confirmed variants among B6J and other inbred mouse strains. Genotyping of strains with high (red) or low (blue) expression of *Gabra2* revealed that that the likely causal variant is the intronic indel (single base pair deletion in B6J). (B) Evidence for disruption of mRNA processing is shown as an accumulation of intronic reads in the affected and downstream introns in B6J (blue) compared to D2 (red). Normalized and binned read pileup of poly-A enriched striatal RNA-seq data from 10 to 11 B6 and D2 individuals, respectively, generated by Bottomly and colleagues (Bottomly et al., 2011) and hosted for viewing and analysis at the GeneNetwork mirror of the UTHSC Genome Browser (http://ucscbrowser.genenetwork.org/). *Gabra2* gene model is shown above (5’ to 3’ orientation with dashed line representing introns and solid vertical bars representing exons). (C) Average hippocampal expression of *Gabra2* exons and introns based on inheritance of B6J (B) and D2 (D) alleles in the BXD population. All coding exons and UTRs showed significantly higher expression associated with inheritance of the D allele. In contrast, introns 4, 5, and 6 showed significantly higher expression associated with inheritance of the B alelle. The variant is located in intron 4. Introns 3, 6 (distal to the fifth coding exon), 7, and 8 are not significantly different based on expression of parental alleles. There was no evidence of a splicing defect resulting in loss of exons in B6J. GeneNetwork data set GN206, UMUTAffy Hippocampus Exon (Feb09) RMA was used for the analysis and includes 45 BXD strains from BXD33 and above. 15 strains inherited the B allele and 30 inherited the D allele at the *Gabra2* locus.

### Proteomics analysis by 6-plex TMT-LC/LC-MS/MS

The analysis was performed essentially based on the previously reported protocol (Bai et al., 2017; Niu et al., 2017). In brief, hippocampus was dissected from male B6J and D2 mice to create pooled samples consisting of three animals each. Pooled samples represented B6J mice aged approximately 60 days (two pooled samples) or 1 year (one pooled sample) and D2 mice aged 90 days (two pooled samples) or 1 year (one pooled sample). Tissue from pooled samples was lysed and digested into peptides. After desalting, the peptides were labeled with TMT reagents and equally mixed. The labeled samples were further fractionated by neutral pH reverse phase liquid chromatography. A total of 10 fractions were collected and further analyzed by low pH reverse phase liquid chromatography (LC), and each fraction was analyzed by acidic pH reverse phase LC-MS/MS.

Collected data was searched against a database to identify peptides and filtered to achieve a 1% protein false discovery rate. TMT intensities were extracted, filtered, normalized, and summarized into peptide and protein quantification results. A total of 22,005 proteins representing 7,074 protein groups were identified and 7,014 protein groups were quantified. Statistical analysis was performed to determine cutoff for altered proteins and to evaluate associated false discovery rate. In this case the significance threshold was set at *p* < 0.005 with a minimum of 3 peptides detected. A total of 8 peptides and 34 spectral counts were detected for *Gabra2*.

### Generation of CRISPR-engineered mice

Knockin mice (B6-*Gabra2*^em1Geh/^J) were produced with CRISPR/*Cas9* using techniques as described (Blednov et al., 2017). Briefly, a sgRNA targeting *Gabra2* at the intron/exon junction near Chr 5 at 71,014,638 Mb (GRCm38/mm10 assembly) was identified using the WTSI Genome Editing database (Hodgkins et al., 2015). Two partially overlapping PCR primers (F:GAAATTAATACGACTCACTATA<u>GGAATTGTAAATTTATATTT</u>GTTTTAGA GCTAGAAATAGC; R:AAAAGCACCGACTCGGTGCCACTTTTTCAAGTTGATAACGGACT AGCCTTATTTTAACTTGCTATTTCTAGCTCTAAAAC) were used to generate a T7 promoter containing sgRNA template (sgRNA target sequence underlined in above sequence) as described (Bassett et al., 2014). The sgRNA and Cas9 mRNA were produced by in vitro transcription, purified using the MEGAclear Kit (Ambion), ethanol precipitated, and re-suspended in DEPC treated water. A 121 nucleotide single stranded DNA repair template oligo (TTTATAGGCTTACTACTTCTAAAACATGTACTGTTTTCAAAGGAATTGTAAATTTAT ATTT***T***AGGAGTATACAATAGATGTTTTCTTTCGGCAAAAATGGAAAGATGAGCGTTTAAAATTT) harboring the desired T insertion (bold, italics in above sequence) in the intron of *Gabra2* was purchased as Ultramer DNA (Integrated DNA Technologies, Coralville, IA). sgRNA (100 ng/µl), *Cas9* mRNA (75 ng/µl), and repair oligo (100 ng/µl) were injected into the cytoplasm of B6J one-cell embryos as described (Yang et al., 2014; Blednov et al., 2017). Pups resulting from injected embryos were screened for DNA sequence changes in intron 3 of the *Gabra2* gene by PCR/DNA sequence analysis. Briefly, a 447 bp amplicon spanning the knockin site was PCR amplified with forward (CAACCAGGAGGGGAAAGACA) and reverse (TTCGAAGCAGCTTGTTTGGC) primers. PCR products were sequenced directly or subcloned into pCR2.1-TOPO (Invitrogen) and sequenced. All mice were genotyped by DNA sequence analysis (Sanger sequencing) using this approach.

The male founder (F0) was subsequently crossed to a female B6J mouse to generate F1 progeny. F1 mice were crossed to generate F2 mice. The colony is maintained through heterozygous breeding and all molecular and behavioral phenotyping were performed in generations F2 and higher.

### Analysis of CRISPR/Cas9 off-target effects

The CRISPOR program (CRISPOR v4.0; crispor.tefor.net) was used to search for off-target sites (Haeussler et al., 2016). For precise genome editing, the sgRNA must match a 20 nucleotide target sequence in the genomic DNA (protospacer sequence) that includes a protospacer adjacent motif (PAM) with sequence NGG (where N is any nucleotide). The sgRNA target sequence (TGAATCATAAACTTATATTTAGG) and PAM used to generate CRISPR-engineered mice was predicted to generate 245 off-targets using the CRISPOR tool. However, all predicted off-targets included two (8 predicted off-target sites), three (57 predicted off-target sites), or four (380 predicted off-target sites) mismatches in the 12 bp adjacent to the PAM site. Perfect matches are more likely to generate off-target effects compared to mismatches. All off-targets were given an off-target score (cutting frequency determination or CFD score (Doench et al., 2016)) in the CRISPOR program. We ranked candidates by the CFD score and selected the top 15 predicted off-target sites for confirmation using traditional capillary sequencing (described above). All of these sites contained at least 4 mismatches and were located in intergenic or intronic regions (STable 1). All sites were located in intergenic or intronic regions with high sequence complexity and repetitive sequences. As a result, two of the primer pairs failed to produce a PCR product, even when using B6J genomic DNA.

### Analysis of global gene expression in CRISPR-engineered Gabra2 mice

Cortex, hippocampus, and striatum were harvested from three male and three female Gabra2^-/-^ and Gabra2^KI/KI^ mice aged 100 to 130 days-of-age as described above. RNA was extracted using the RNeasy Lipid Tissue kit (Qiagen) and contaminating genomic DNA was removed following treatment with DNAse (Qiagen). RNA quantity was measured using a Nanodrop2000 Spectrophotometer (ThermoScientific) and quality was assessed using the RNA nano chip and Bioanalyzer (Agilent). All samples had RNA integrity values (RIN) over 8. Gene expression was measured using the Clariom D Assay mouse (Thermo Fisher Scientific). Sample preparation and hybridization was performed according to the manufacturers protocol and performed by the UTHSC MRC Institutional Core. Affymetrix Expression Console Software (Thermo Fisher Scientific) was used to annotate and process raw probe cell intensity (CEL) files. The Gene Level-SST-RMA algorithm was used to perform signal space transformation (SST) and guanine cytosine count correction prior to robust multi-array average (RMA) normalization on the batch of 36 samples (3 brain regions × 2 sexes × 3 biological replicates × 2 genotypes). There were no outliers following normalization. The data set was filtered to include only “Main” category probe sets corresponding to annotated coding and non-coding transcripts with a variance > 0.1 across all samples. This reduced the data set from 72,688 to 12,699 probe sets. This data set was used for enrichment analysis, comparison of downstream effects of Gabra2 variation in the BXD family of strains, and to investigate possible compensatory changes in other α subunits following reduction of *Gabra2*. For each analysis, genes with differential expression between Gabra2^KI/KI^ and Gabra2^-/-^ were detected using Students t-test in R to compare expression of each gene by genotype. Male and female samples of each genotype were pooled for the analysis.

Enrichment analysis was performed for differentially expressed genes (*p* < 0.01, *n* = 684) in the striatum using WebGestalt (Zhang et al., 2005; Wang et al., 2013). Overrepresentation of functional terms in the differentially expressed gene set compared to the reference set (consisting of the 12,699 filtered probe sets) was calculated using the Overrepresentation Enrichment Analysis (ORA) method using the following functional databases: Gene Ontology (GO) biological process, GO cellular component, GO molecular function, KEGG, and mammalian phenotype ontology (MPO). Significant enriched terms were identified using the following criteria: (1) minimum number of three genes in a category and (2) a False Discovery Rate (FDR) of > 0.05 or nominally significant *p*-value < 0.01. For analysis of compensatory changes, the expression of all major subunits was compared between genotypes in the cortex, striatum, and hippocampus.

## Results

### Reduced levels of Gabra2 occur only in the B6J substrain

The B6J strain showed the lowest levels of *Gabra2* among 16 common and wild-derivative inbred strains, including the closely related C57BL/6ByJ substrain (Fig. 1A). This difference was robustly detected across expression platforms and was not dependent on sex or brain region. For example, average log2 expression of *Gabra2* in the hippocampus was 9.3 for B6J and 11.2 for D2 with over 3.6-fold variation across inbred strains of mice (Affymetrix M430 array, Fig. 1A). The decrease in *Gabra2* mRNA levels was global, and this reduction relative to D2 was also detected in 10 mesocorticolimbic regions (RNA-seq, Fig. 1B). To evaluate whether reduced levels of *Gabra2* were only evident in B6J, we examined expression in a small pedigree of C57BL/6 substrains with defined dates of separation—B6J, B6(Cg)-TyrC-2J/J(B6C2J), C57BL/6EiJ (B6EiJ), C57BL/6JByJ (B6ByJ), C57BL/6NJ (B6NJ), and C57BL/6NCrl (B6NCrl) (Fig. 1C). Of these substrains, B6EiJ was the most recently separated from B6J in 1976. The albino strain (B6C2J) was maintained as an inbred colony since the discovery of the albino mutation around 1970 (Townsend et al., 1981) but over the past decade has been repeatedly backcrossed to B6J stock and is therefore now listed as a congenic (Cg) strain of B6J rather than as an independent substrain. Expression of *Gabra2* in cortex, striatum, and hippocampus was significantly higher (p < 0.05) in B6EiJ, B6ByJ, B6NJ, B6NCrl, and D2 as compared to B6J and B6C2J (Fig. 1D). These results explain the remarkable variation in *Gabra2* levels (over 4-fold) observed both within sets of BXD strains based on time of generation between the late 1970’s and the early 2000’s (Taylor et al., 1999; Peirce et al., 2004) (Fig. 1D–1E, SFig3) and recent crosses between B6J and C3H/HeJ or B6J and CAST/EiJ (Mulligan et al., 2012). Lower expression in these crosses was always associated with inheritance of the B6J parental allele of *Gabra2*. F2 intercrosses can also be used to estimate the effect of each allele on expression and mode of inheritance. In F2 crosses in which B6J is a parental strain, the B6J allele was completely recessive (SFig4). Taken together, there is strong evidence that a spontaneous mutation in the *Gabra2* gene was fixed in the B6J foundation stock colony at the Jackson Laboratory between 1976 and 1991. This mutation greatly reduces brain expression of *Gabra2* mRNA in B6J and all known derivative strains.

To evaluate whether the private B6J mutation also leads to a reduction in protein level, we measured GABRA2 expression using traditional Western blots and shotgun proteomics methods. We first compared expression between B6J and D2 using an unpublished hippocampal proteome dataset generated using the tandem mass tag (TMT) approach. Significantly (*p* < 0.005) less Gabra2 normalized peptide counts were detected in B6J (0.77 ± 0.04) relative to D2 (1.23 ± 0.07) (Fig. 1G). To confirm and expand upon this finding, we probed with antibodies specific for Gabra2 in the hippocampus and also detected a marked reduction in protein levels in the B6J strain compared to both the D2 strain and the B6NJ substrain (Fig. 1H, SFig 1).

### Identification of candidate non-coding sequence variants modulating Gabra2 expression

The G*abra2* gene resides in a genomic region that is largely identical by descent among inbred strains, with the exception of the wild derived strains (e.g. CAST/EiJ, PWK/PhJ, and SPRET/EiJ) and contains very few known sequence variants, most of which are located within intronic or intergenic regions. B6J and derivative congenic strains, such as B6C2J, are the lowest expressing individuals from a panel of inbred strains and C57BL/6 substrains (Fig. 1). Thus, the causal sequence variant must share a common pattern that differentiates B6J from all other inbred strains and substrains. To identify candidate variants near the *Gabra2* locus we used existing resources generated by our group (100X coverage of the D2 genome available at http://ucscbrowser.genenetwork.org/ (Wang et al., 2016)) and the Wellcome Trust Sanger Institute (>30X average coverage of the genomes of 36 inbred mouse strains, including the B6NJ substrain available at http://www.sanger.ac.uk/cgi-bin/modelorgs/mousegenomes/snps.pl). Using these resources, we identified one SNP and two insertion/deletions (indels) that were of high sequence quality. All three were private to B6J compared to B6NJ and D2. We were able to independently validate one SNP and one indel by capillary-based dideoxynucleotide sequencing. Validated candidate polymorphisms were located either ~30 Kb downstream of the *Gabra2* gene locus in an intergenic region with high conservation among the mammalian lineage (SNP rs29547790 located on Chr 5 at 70.93 Mb) or in an intron (the third nucleotide from exon 4) adjacent to a splice acceptor site (indel located on Chr 5 at 71.01 Mb) (Fig. 2A).

To identify the causal variant, we genotyped D2, C57BL/6 substrains, and BXD strains that differed in the level of *Gabra2* expression based on inheritance of the presumed wild type allele prior to the late 1970s (D2, B6ByJ, B6NJ, B6EiJ and BXD29) or the mutant allele which was fixed in the B6J substrain between 1976 and 1991 (B6J and BXD40). Importantly, only the genotype at the intronic variant was perfectly associated with *Gabra2* expression levels (Fig. 2A). Strains with the mutant allele (single base deletion), such as B6J and BXD40, had low expression of *Gabra2* compared to the five strains with the wild type allele. The intronic variant was located near a splice acceptor site but the specific impact on splicing is not known. However, striatal RNA-seq data (Bottomly et al., 2011) generated for B6J (n = 10) and D2 (n = 11) revealed a higher number of reads mapping both to the intron containing the enigmatic deletion and two downstream introns in the B6J strain (Fig. 2B) which could indicate errors in mRNA processing. Significantly higher intronic expression (probe 4462485; *p* < 0.001) was also detected among BXD strains (BXD33 through 100) that inherited the B6J allele (6.1 ± 0.22) compared to those that inherited the D2 allele (5.06 ± 0.13) using an exon microarray platform (GeneNetwork data set GN206, UMUTAffy Hippocampus Exon (Feb09) RMA; Fig. 2C). All *Gabra2* exons are expressed at the mRNA level in BXD strains regardless of allele, suggesting that the variant does not produce alternative mRNA isoforms. Instead, all coding exons have higher expression in strains that inherited the D2 (D) allele, and three introns containing or immediately downstream of the variant have higher expression in strains that inherited the B6J (B) allele (Fig. 2C). In sum, a non-coding intronic deletion is the sole candidate variant responsible for decreased *Gabra2* mRNA and protein expression in B6J.

### Repair of the candidate non-coding variant is sufficient to restore Gabra2 levels in B6J

To test whether the intronic deletion is sufficient to globally reduce brain mRNA and protein in B6J we used the CRISPR (clustered regularly interspaced short palindromic repeats) and CRISPR associated (*Cas9*) system to repair the mutation on a pure B6J genetic background (*Gabra2*^B6J/B6J^; Fig. 3A). Insertion (knockin; KI) of a single nucleotide was sufficient to fully restore *Gabra2* protein (Fig. 3B, SFig 2) and mRNA levels (Fig. 3C) in brain tissue of B6-*Gabra2*^em1Geh/^J homozygous knockin (KI, *Gabra2*^KI/KI^) and heterozygous (HET, *Gabra2*^KI/B6J^) F2 offspring (Fig. 3D). These results are consistent with the recessive mode of inheritance of the mutant and hypomorphic B6J allele (*Gabra2*^B6J/B6J^) that is associated with reduced expression due to the presence of the intronic deletion. The use of CRISPR engineering can produce both off-target effects and founders that are mosaic for the introduced mutation (*e.g.* some cells may have the mutation, some may not, and other cells may have different mutations). However, the expression level of Gabra2 protein and mRNA in the brain tissue of the founder mouse was equivalent to strains without the Gabra2 deletion (*e.g.* ByJ and NJ; Fig. 3B and 3C) indicating correction in most, if not all, neurons of the founder mouse. In addition, no off-target modifications were detected in the top 15 predicted off-target sites in the F0 mouse and his progeny compared to the B6J reference genome (see methods and STable 1).

**Fig. 3.**
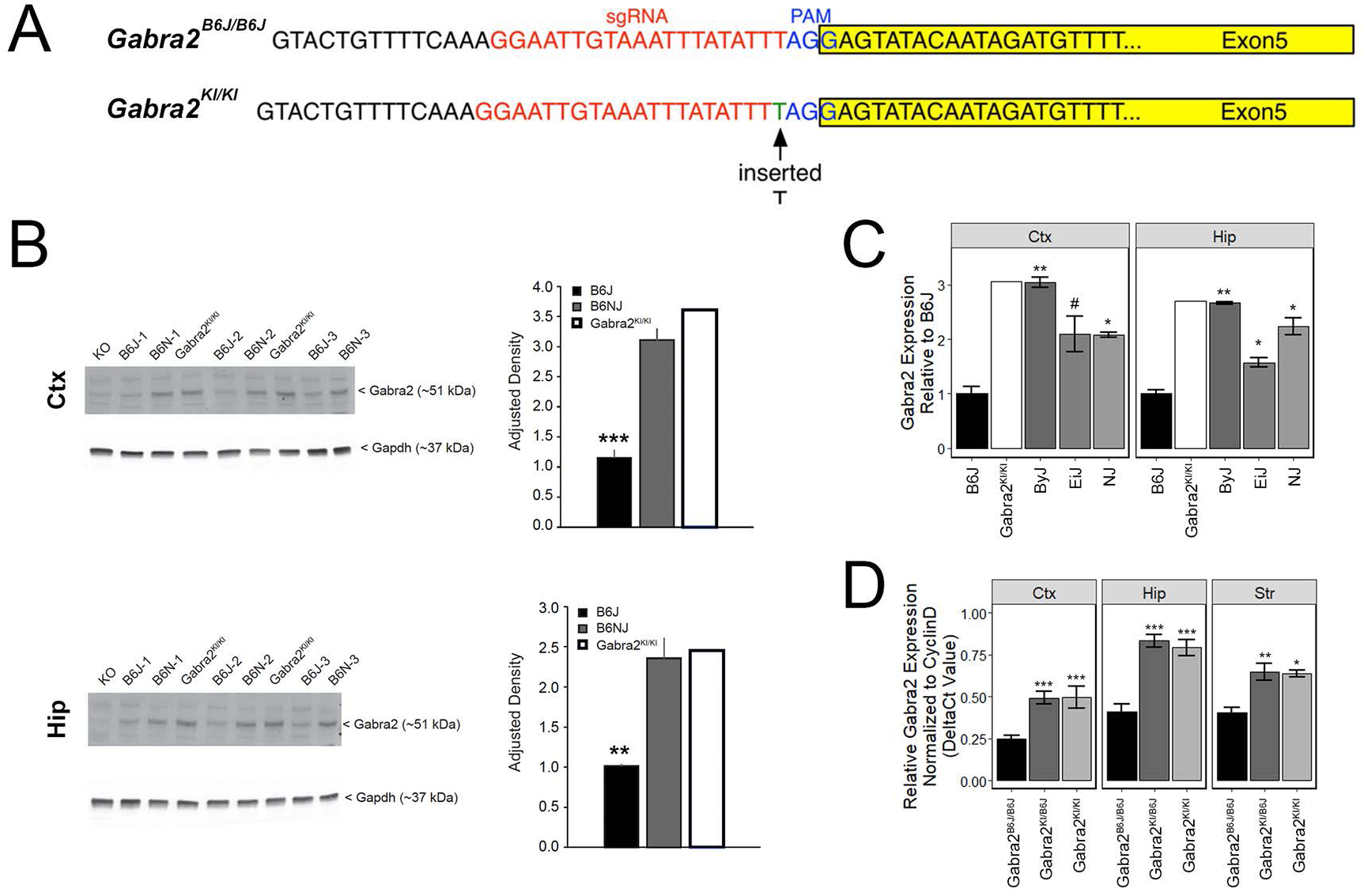
Repair of private deletion in B6J is sufficient to restore *Gabra2* mRNA and protein levels. (A) Site of repair of the private deletion in B6J (green), guide RNA (red) and PAM site (blue) is shown. The exon is highlighted in yellow. (B) Protein levels of *Gabra2* are also restored in the hippocampus (top panel) and cortex (bottom panel) of the original Gabra2^KI/KI^ founder mouse (n = 1) relative to B6J (n = 3) and are similar to that of B6NJ (n = 3). Western blot shown to the left and adjusted density bargraph shown to the right for each region. (C) Hippocampal and cortical *Gabra2* mRNA levels were restored in the original Gabra2^KI/KI^ founder after insertion of a single nucleotide relative to B6J. Expression of *Gabra2* in the founder mouse was similar to that of other B6 substrains. Expression was measured viaqPCR and shown relative to B6J. Males and females were combined for the statistical analaysis (n = 5 to 6 per strain). Note that many CRISPR founders were mosaic for the introduced mutation such that some cells may have the mutation, some may not, and other cells may have different mutations. We include molecular validation in the founder here in order to demonstrate that our engineering approach worked, and that mosaicism is not evident in brain tissue of the Gabra2^KI/KI^ founder mouse. Tissue from *Gabra2* knockout mice (KO, predominately B6J background mixed with 129/SvEv, see methods) was kindly provided by Y. Blednov from The University of Texas and used as a negative control. (D) Hippocampal, striatal, and cortical *Gabra2* mRNA levels were also restored in heterozgous Gabra2^KI/-^ and homozygous knockin Gabra2^KI/KI^ mice after insertion of a single nucleotide relative to homozygous Gabra2^B6J/B6J^ mice. Expression measured via qPCR and data were analyzed at the ΔCt level (relative to the expression of the control gene CyclinD). There is a significant main effect of genotype [F(2,32) = 47.1, p = 2.9e-10] and brain region [F(2,32) = 27.89, p = 9.7e-08]. The interaction was not significant. Pairwise significance between genotypes and within brain region were determined using Tukey’s HSD test and shown relative to control B6J or Gabra2^B6J/B6J^ mice. Significance defined as: # = *p* < 0.1, * = *p* < 0.05, ** = *p* < 0.01, *** = *p* < 0.001.

### Intronic Gabra2 deletion is associated with alterations in GABAergic striatal gene expression and signaling

To precisely identify biological processes or pathways altered by variation in *Gabra2* levels in the striatum, hippocampus, and cortex, we performed enrichment analysis on genes differentially expressed (*p* < 0.01; 684 genes) between homozygous Gabra2^KI/KI^ mice with “normal” *Gabra2* levels and "mutant" Gabra2^B6J/B6J^ mice with low levels of *Gabra2*. No significantly enriched categories were observed for differentially expressed genes from the cortex or hippocampus. However, a number of enriched categories (FDR adjusted p-value < 0.05) related to GABAergic synaptic signaling were detected in striatum, including ion transmembrane transport, localization to the synapse, GABA receptor activity, retrograde endocannabinoid receptor signaling, dopaminergic synapse, cAMP signaling pathway, and GABAergic synapse (STable 2 and STable 3). Mammalian Phenotype Ontology terms related to emotional behavior, cognition, anxiety, sensorimotor gating, fear, and reward were also enriched, albeit at a nominally significant level. These results indicated that alterations in GABAergic signaling, specifically in the striatum, might be a direct downstream consequence of reduced *Gabra2* levels.

Alterations in GABAergic signaling could result from the loss of Gabra2-containing receptors or changes in the composition of GABA-A receptors to reflect inclusion of other alpha subunits in the absence of Gabra2. To begin to address these issues, we compared the transcript levels of all major GABA-A receptor subunits (α1-6; β1-3, γ1-3, and δ) in multiple brain regions between homozygous *Gabra2*^KI/KI^ mice with normal *Gabra2* levels and mice harboring the mutant B6J allele (*Gabra2*^B6J/B6J^). In addition to a large and significant reduction in *Gabra2* mRNA, the transcript levels of several major GABA-A receptor alpha subunits (*Gabra1*, *Gabra3*, *Gabra5*) were also significantly reduced in striatum of *Gabra2*^B6J/B6J^ mice relative to the *Gabra2*^KI/KI^ mice (Fig. 4). In contrast, the level of *Gabra4* was modestly increased in *Gabra2*^B6J/B6J^ mice. In striatum, alterations in alpha subunits were accompanied by significant decreases in beta (1 and 2) and gamma (1 and 2) subunits and a significant increase in the delta subunit (B6J allele relative to KI allele; Fig. 4). Highly significant alterations in alpha subunits were not observed in the hippocampus or cortex between genotypes, but there was a trend for increased expression of both the α1 subunit in the cortex and the α3 subunit in the hippocampus (B6J allele relative to KI allele). In the cortex there was an associated trend for increased expression of the β1 subunit and decreased expression of the γ1 subunit. In the hippocampus, there was a significant decrease in γ1 subunit levels and a trend for decreased expression of the β1 and γ3 subunits. Taken together, these results suggest that decreased levels of *Gabra2* associated with the B6J allele are also associated with profound alterations in the level and composition of GABA-A receptor subunits, especially in the striatum.

**Figure 4.**
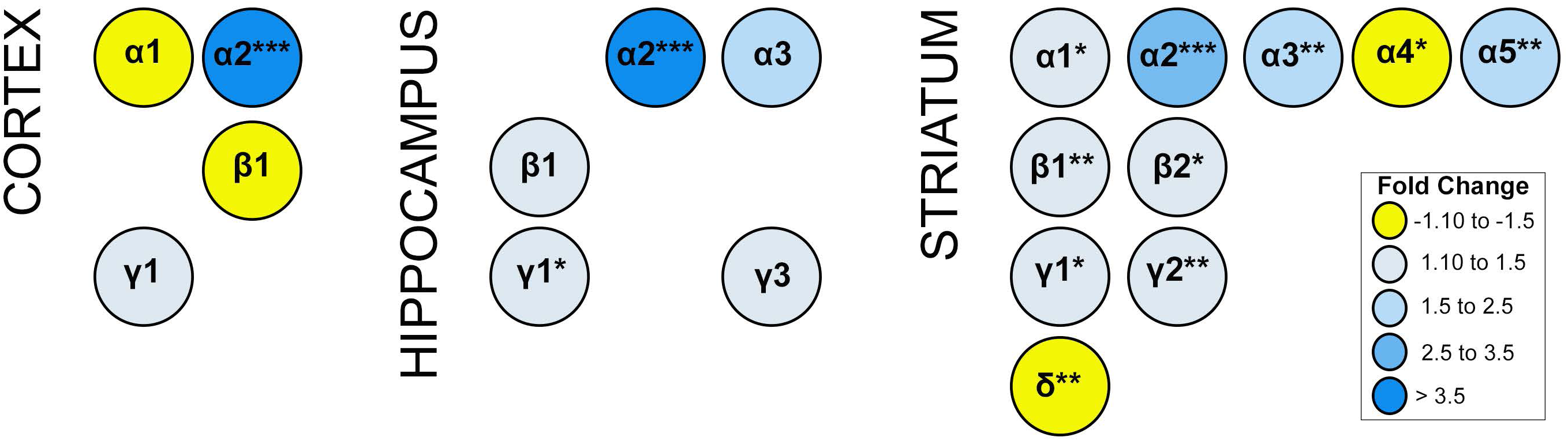
Expression of GABA-A Receptor Alpha Subunits. Expression generated using the Affymetrix Clariom D Assay (microarray platform). Only subunits with significant or suggestive (p < 0.1) differential expression between B6J and KI *Gabra2* genotypes are shown. Signifcance defined as: * = *p* < 0.05, ** = *p* < 0.01, *** = *p* < 0.001. Fold change is indicated by color intensity with yellow representing increased expression in Gabra2^B6J/B6J^ (B6J allele) mice relative to Gabra2^KI/KI^ (KI allele) mice. In contrast, blue represents decreased expression in B6J allele mice relative to KI allele mice. Alterations in several alpha subunits, inlcuding the major *Gabra1* subunit, are observed in the cortex and striatum of B6J allele mice which harbor a naturally occuring intronic deletion resulting in decreased *Gabra2* expression. The only alpha subunit with significant and higher expression (fold change > 1.3) in B6J allele mice is *Gabra4* (striatum). Beta and gamma subunits are also altered at a suggestive level in cortex and hippocampus. In the hippocampus, γ1 expression is significantly reduced in mice with the B6J allele. In the striatum, β1-2 and γ1-γ2 are also significantly reduced in mice with the B6J allele of *Gabra2*. In contrast, striatal levels of the delta subunit are increased in mice with the B6J allele relative to those with the KI allele. These results suggest brain region specific alterations in the abundance of certain classes of GABA-A receptors associated with inheritance of the *Gabra2* B6J allele. For example, in the cortex, α1β1-containing receptors may be increased while α2γ1-containing subunits are decreased. In the hippocampus α2 and α3-containing receptors including β1 or γ1-γ2 subunits may be decreased. In the striatum there is evidence that there may be a general decrease in receptors containing α1, α2, α3 or α5; and β1 or 2; and γ1 or γ2. In contrast, there may be an increase in receptors containing α4 and δ subunits in striatum.

## Discussion

Using a combination of genetic, genomic, functional, and bioinformatics approaches we have identified a private and non-coding single nucleotide deletion in the *Gabra2* gene in the B6J strain that results in substantial reduction of brain mRNA and protein levels (Fig. 1, Fig. 2). Repair of this deletion using CRISPR-*Cas9* genome editing fully restored *Gabra2* levels (Fig. 3) on the B6J genetic background. The resulting genetically engineered rodent model system (*Gabra2*^B6J/B6J^ and *Gabra2*^KI/KI^ mice) is thus, a valuable resource for probing the molecular and behavioral consequences of alterations in *Gabra2* expression. These results have important implications for any investigators using B6J as their background strain for molecular genetic studies of neurobiology and behavior, in particular those involving study of the GABA system.

Unlike previous studies in which function of *Gabra2* was probed through complete genetic deletion, the mutation associated with the B6J allele of *Gabra2* is hypomorphic and not a complete null. Complete deletion of function represents an extreme case in biological systems and there are often many compensatory changes that can make interpretation of results difficult. Compensation by other α subunits has not been profiled extensively in *Gabra2* KO mouse lines and some differences in behavior could be due to enhanced expression of other subunits and differences in genetic background as existing *Gabra2* KO mice differ in the proportion of B6J and 129/SvEv genetic background. In contrast, our study probed the role of natural variation in the level of *Gabra2* in the context of an isogenic B6J background and profiled the expression of all major GABA-A subunits in cortex, striatum, and hippocampus—regions involved in different aspects of mood disorders and addiction. Surprisingly, even in the context of an isogenic B6J background, we detected many alterations in GABAergic signaling at the transcriptional level and observed changes in GABA-A receptor subunit expression between *Gabra2*^B6J/B6J^ and *Gabra2*^KI/KI^ in all three regions with the most profound alterations detected in the striatum. Regardless of brain region, the major alteration (fold change >2.5) was the loss of *Gabra2* subunits. However, the levels of other alpha subunits were also altered (Fig. 4), albeit at a much lower level (<1.5 fold). Exceptions include the levels of *Gabra5* (1.7 fold decrease relative to *Gabra2^KI^*^/KI^) and *Gabra3* (2.5 fold decrease relative to *Gabra2*^KI/KI^) in the striatum. There are no known mutations in *Gabra3* or *Gabra5* in the B6J background, so the reduction in their levels in striatum must be a direct result of the reduction in *Gabra2*. Based on the available data, our hypothesis is that there are simply fewer α2 containing GABA-A receptors in B6J derived strains, and possibly fewer GABA-A receptors in some regions. This is supported by the trend towards decreased cortical α2:γ1 subunit levels, decreased hippocampal α2:β1:γ1 and α2:β1:γ3 subunit levels, and decreases in multiple alpha, beta, and gamma subunit levels in the striatum of *Gabra2*^B6J/B6J^ relative to *Gabra2*^KI/KI^. The pronounced impact of the B6J mutation on GABA-A receptor subunit expression in the striatum, combined with the involvement of the striatum in anxiety and behavioral sensitization to alcohol and other drugs of abuse, strongly suggests that alterations in the level of *Gabra2* in this region drive related behavioral differences between mice with the low-expressing B6J allele and the high-expressing wild type allele. Future functional studies exploring behavior and inhibitory neurotransmission across multiple brain regions in B6-*Gabra2*^em1Geh/^J genotypes will test this hypothesis.

Overall, the findings of our study have broad implications for both mouse and human research. Despite numerous associations between human *GABRA2* variants and addiction-related traits, the mechanisms underlying these associations are unclear. To date, there is no direct link between variants in *GABRA2* and alterations in gene expression or function in human brain. However, if functional variants in *GABRA2* exist, they are almost certainly non-coding variants that exert functional impact through alteration of splicing, gene regulation, or mRNA stability as opposed to variants resulting in a complete loss of function. This may be similar to the function of the non-coding variant controlling variation in *Gabra2* levels in crosses involving B6J and in our B6-*Gabra2*^em1Geh/^J preclinical model system and suggests that non-coding variants in *GABRA2* in human populations might be functional and causal. Evaluating and confirming the functional impact of these variants in human populations is extremely difficult due to the complexity of human genetics and environmental factors as well as the limited ability to assay brain gene expression and circuitry. Therefore, preclinical models such as ours are of vital importance for generating new hypotheses, endophenotypes, and underlying molecular mechanisms to test in human populations. This might include the generation of humanized mice in which candidate functional GABRA2 variants are introduced into the Gabra2 corrected B6J genetic background that is permissive for detection of variants that modulate Gabra2 expression.

We discovered a functional variant causing reduced *Gabra2* expression in B6J, the gold standard reference genome in mouse genetics research and the most widely used inbred strain in biomedical research. It is now critical to re-evaluate gene deletion and other studies in light of the naturally occurring mutation in *Gabra2*. For example, the *Gabra2* locus was recently posited to be a genetic modifier of genetic deletion of *Scn1a*, a voltage-gated sodium channel gene implicated in a spectrum of seizure-related disorders in humans (Hawkins et al., 2016). Hawkins and colleagues observed that the genetic background of *Scn1a*^+/-^ mutations impacted the severity of the epilepsy phenotype such that mutations generated on a B6J background had increased seizures and premature death compared to the same mutation on a 129S6/SvEvTac background, which had a normal phenotype. Our study suggests a specific hypothesis, namely that reduced Gabra2 expression and alterations in GABAergic signaling comprise the causal mechanism underlying enhanced seizure susceptibility induced by *Scn1a* deletion on the permissive B6J background. Likewise, because the B6J strain is also the most widely used reference strain for biomedical research, several findings in this strain may be confounded by alterations in *Gabra2* and other GABA-A subunits. However, as long as investigators are aware of gene variants with a large impact on expression and function, such as the non-coding variant in *Gabra2*, these naturally occurring mutations can be vital tools for discovery and systems genetics (Li et al., 2010; Mulligan and Williams, 2015; Wang et al., 2016). The CRISPR-*Cas9* B6J *Gabra2* repaired line (B6-*Gabra2*^em1Geh/^J) will serve as a crucial resource for directly testing the role of *Gabra2* variation related to phenotypic differences in B6J derived lines and substrains.

## Acknowledgments

This research was partially supported by NIAAA INIA grants U01AA13499 and U01AA016662 to Drs. Mulligan, Lu, and Williams; NIH grant R01AG047928 to Drs. Peng and Wang; NIH grants R21AG048446 to Dr. Kaczorowski and F31AG050357 to S. Neuner; NIDA grants R03DA038287 and R21DA038738 to Dr. Bryant; and NIAAA grants AA010422 and AA020889 to Dr. Homanics. We also thank Carolyn Ferguson (University of Pittsburgh), Casey Bohl (University of Tennessee), and Melinda McCarty (University of Tennessee) for superb technical support.

---

**STable 1. Summary of Predicted Off-Targets.** Predictions made using the CRISPOR program and prioritized based on CFD score.

**STable 2. Overrepresented functional terms for differentially expressed genes in the striatum of CRISPR engineered *Gabra2* KI mice.** Gene members are shown for categories with less than 20 members. All gene members for each category are shown in STable 3.

**STable 3. Summary of genes with significantly enriched functional terms.** Summary of results are provided in the first worksheet (C = total number of genes in category; O = number of observed genes in category; E = number of genes in category expected by chance). Remaining worksheets contain all enriched categories and genes.

**Supplementary Figure 1:**
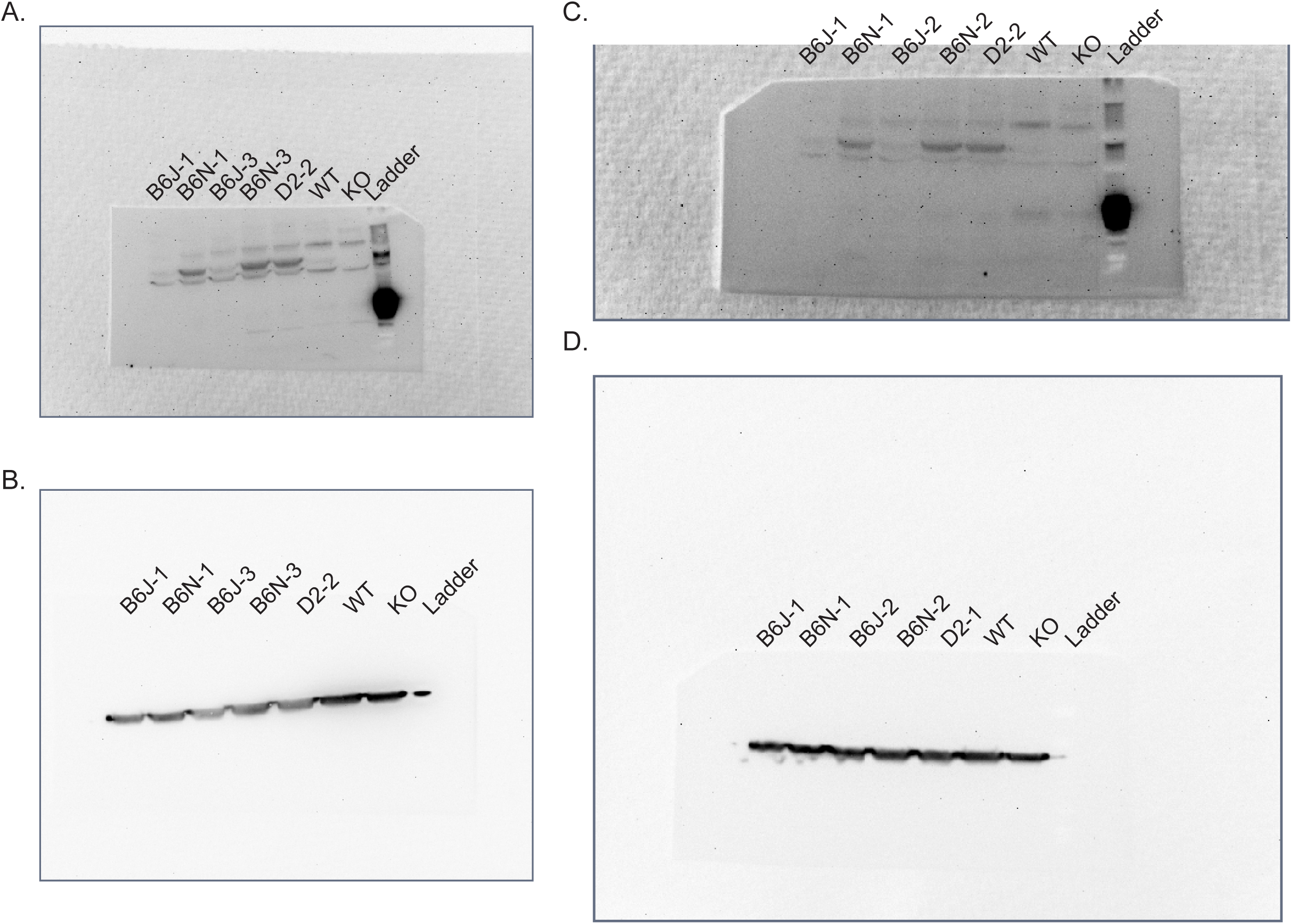
Original blots from founder mouse analysis. Blots were incubated overnight with anti-GABRA2 (PhosphoSolutions #822-GA2CL) and anti-GAPDH (Fitzgerald #10R-G109A) antibodies, followed by fluorescent-conjugated secondary antibodies, and developed on an Odyssey imaging system. (A and B) show unedited whole-blot GABRA2 and GAPDH staining of cortex samples, respectively. (C and D) show corresponding GABRA2 and GAPDH uncropped whole-blot images that have been edited using Photoshop’s brightness/contrast and levels features to make bands visible. (E-H) show equivalent images for blots containing hippocampal samples.

**Supplementary Figure 2.**
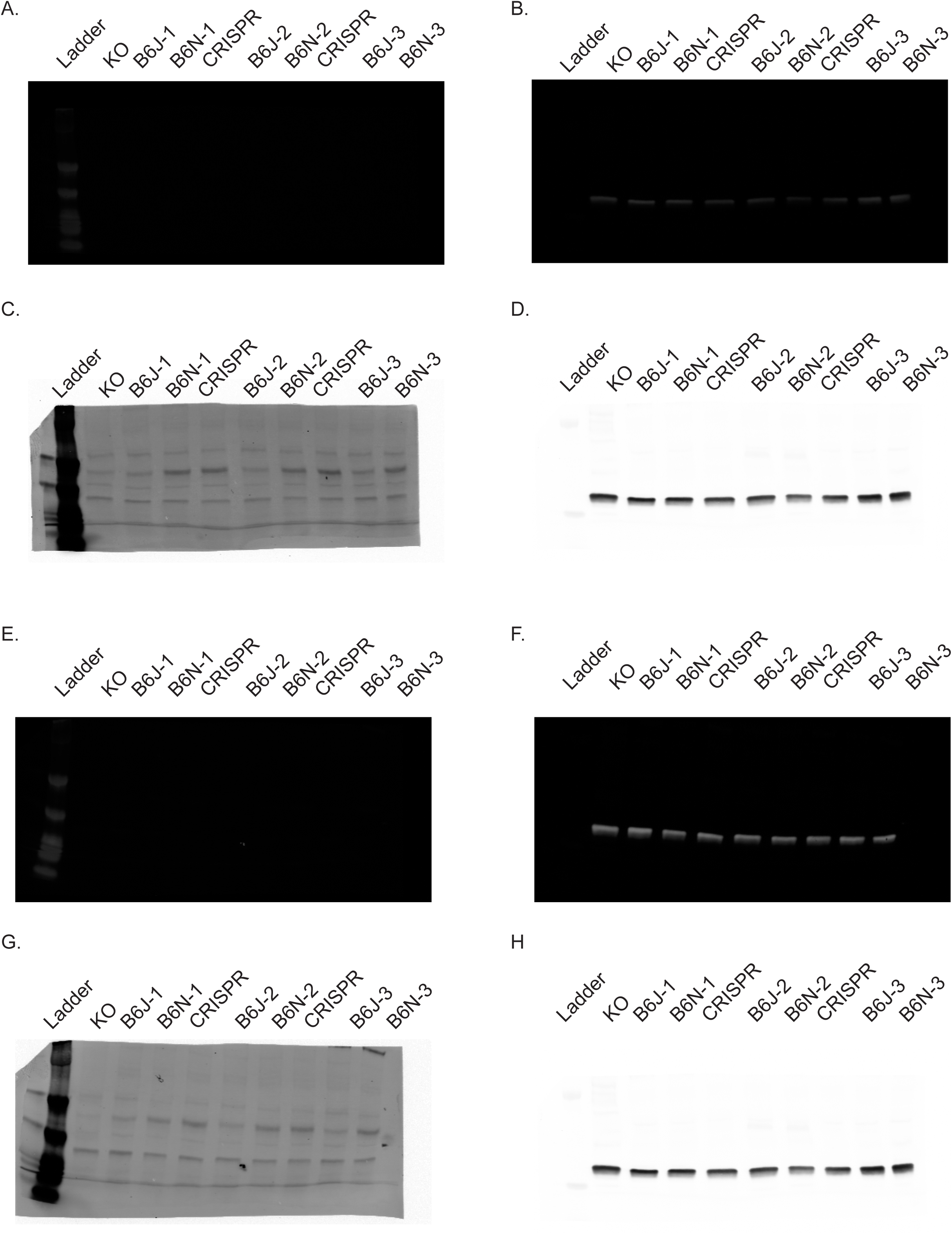
Original blots from Figure 4 analysis. Blots were incubated overnight with anti-GABRA2 antibody (PhosphoSolutions #822-GA2CL), followed by a horseradish peroxidase-conjugated anti-rabbit secondary, and developed on a BioRad ChemiChem chemiluminescent detection system. (A) Unedited whole-blot GABRA2 staining. Following development, blots were stripped and re-probed for GAPDH (Fitzgerald #10R-G109A) as a loading control, and developed using the same protocol. (B) Unedited whole-blot GAPDH staining. (C and D) show equivalent whole-blot unedited images from a second replicate of the experiment. Samples B6J-1 and B6N-1 were run during both experiments.

**Supplementary Figure 3.**
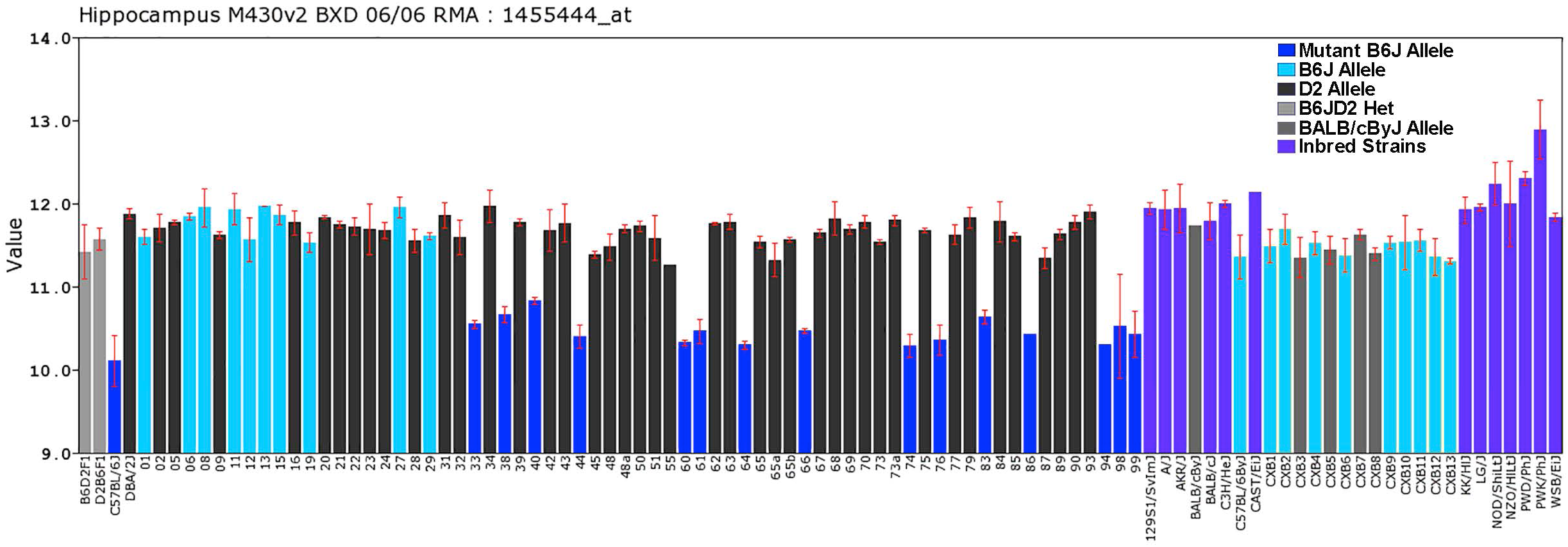
Strain distribution pattern of expression of *Gabra2*. A 2 to 3-fold reduction in expression of mRNA levels in hippocampus [GN110 Hippocampus Consortium M430v2 (Jun06) RMA] that is only segregating in the new BXD strains (BXD33 and higher) that have inherited the *Gabra2* B6J private mutation. The Y-axis provides an estimate of the expression of *Gabra2* (Affymetrix probe set 1455444_at) on log_2_ scale. To the far left are the two reciprocal F1 hybrids between B6J and D2 with comparatively normal high expression, demonstrating a dominant phenotype of the wild type D allele. Note that all of the initial set of BXDs (from BXD1 to BXD32) have high expression, a finding only compatible with the hypothesis that the mutation in B6J occurred after the inception of these strains. In contrast, 15 of the newer BXD strains have low expression. Finally, all other common inbred strains, including both B6ByJ and all parental strains of the Collaborative Cross other than B6J have high expression. High expression is also evident in the CXB recombinant inbred panel generated by crossing BALB/cByJ and B6ByJ. Bars colored by genotype or inbred strain. BXD genotype determined using marker rs13478320 located on Chr 5 a 70.742059 Mb. CXB genotype determined using marker rs13478300 located on Chr 5 at 65.043224 Mb.

**Supplementary Figure 4.**
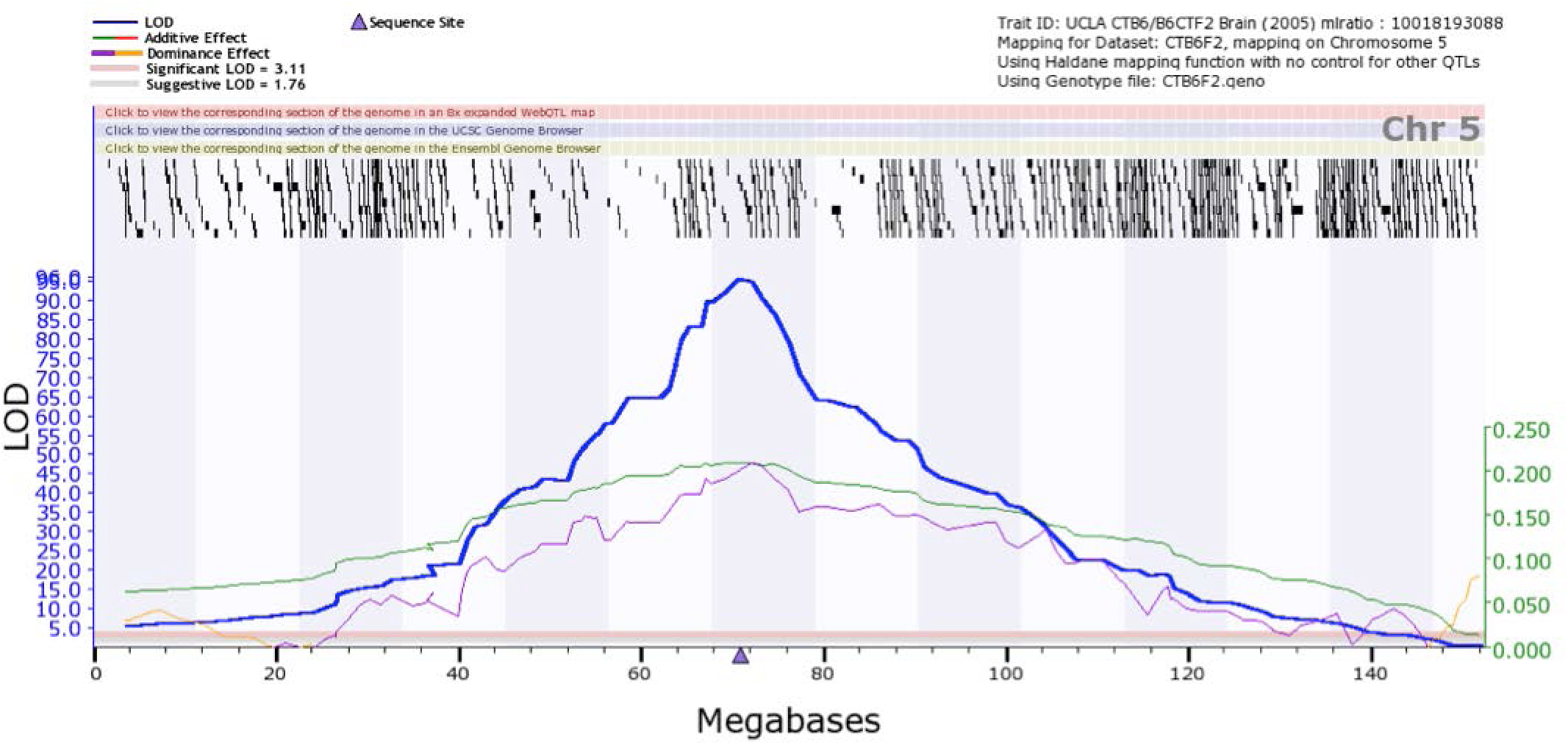
Mode of inheritance of *Gabra2* alleles. In a large eQTL transcriptome analysis of 400 F2 intercross progeny between B6J and CAST/EiJ, the LOD peak is precisely aligned on the *Gabra2* gene (purple triangle), the effect size is about 0.20 z per allele (right Y-axis). Note also that the dominance effect is complete (compare peak of the purple and green effect size plots).

